# The neurophysiological basis of short- and long-term ventriloquism aftereffects

**DOI:** 10.1101/2020.06.16.154161

**Authors:** Hame Park, Christoph Kayser

## Abstract

Our senses often receive conflicting multisensory information, which our brain reconciles by adaptive recalibration. A classic example is the ventriloquist aftereffect, which emerges following both long-term and trial-wise exposure to spatially discrepant multisensory stimuli. Still, it remains debated whether the behavioral biases observed following short- and long-term exposure arise from largely the same or rather distinct neural origins, and hence reflect the same or distinct mechanisms. We address this question by probing EEG recordings for physiological processes predictive of the single-trial ventriloquism biases following the exposure to spatially offset audio-visual stimuli. Our results support the hypothesis that both short- and long-term aftereffects are mediated by common neurophysiological correlates, which likely arise from sensory and parietal regions involved in multisensory inference and memory, while prolonged exposure to consistent discrepancies additionally recruits prefrontal regions. These results posit a central role of parietal regions in mediating multisensory spatial recalibration and suggest that frontal regions contribute to increasing the behavioral bias when the perceived sensory discrepancy is consistent and persistent over time.

## INTRODUCTION

Sensory recalibration serves to continuously adapt perception to discrepancies in our environment, such as the apparent displacement of the sight and sound of an object (Chen and Vroomen 2013; De Gelder and Bertelson 2003). Despite the importance of such adaptive multisensory processes in everyday life, their neural underpinnings remain unclear. Our environment changes on multiple timescales and not surprisingly, perceptual recalibration emerges also on distinct scales (Bosen et al. 2017, 2018; Bruns and Röder 2015, 2019; Rohlf et al. 2020; Van der Burg, Alais, and Cass 2015). During the ventriloquism aftereffect (Bruns and Röder 2015; Canon 1970; Radeau and Bertelson 1974; Recanzone 1998; Wozny and Shams 2011) the exposure to displaced acoustic and visual stimuli reliably biases the perceived location of subsequent unisensory sounds (Frissen, Vroomen, and de Gelder 2012; Mendonça et al. 2015; Watson et al. 2019; Woods and Recanzone 2004). This aftereffect increases with prolonged exposure to a consistent discrepancy, but independently emerges trial-by-trial and following long-term exposure (Bruns and Röder 2015; Frissen et al. 2012; Kramer, Röder, and Bruns 2020; Van der Burg et al. 2015; Watson et al. 2019). The short- and long-term biases differ in their specificity to the sensory features of the inducing stimuli and hence supposedly constitute two separate processes with distinct neurophysiological correlates (Bruns and Röder 2015, 2019). Still, this hypothesis has not been directly tested.

In a previous study on the ventriloquism aftereffect we showed that medial parietal regions integrate audio-visual information within a trial and mediate the trial-by-trial aftereffect (Park and Kayser 2019), implying a role of parietal regions involved in spatial working memory (Martinkauppi 2000) and multisensory causal inference (Rohe, Ehlis, and Noppeney 2019; Rohe and Noppeney 2015) in short-term recalibration (Wozny and Shams 2011). Given that long-term recalibration results from the prolonged exposure to consistent sensory discrepancies, one could reason that these parietal regions also mediate the long-term effect. However, the few existing neuroimaging studies reported correlates in early sound-evoked potentials and near early auditory cortices (Bruns, Liebnau, and Röder 2011; Zierul et al. 2017), and concluded that the long-term aftereffect is implemented by early sensory regions, in line with evidence from single cell recordings (Recanzone 1998; Recanzone et al. 2000). However, these studies used neural signatures of sound encoding to experimentally test for neural correlates, thus possibly biasing these studies towards auditory pathways. Indeed, one study also reported changes in functional coupling between auditory and parietal regions (Zierul et al. 2017), hinting at a more extensive cerebral network shaping the long-term effect.

We set out to directly compare the neural underpinnings of audio-visual spatial recalibration on a trial-by-trial level (short-term: ST) and after long-term exposure (LT) in human participants. We focused on the hypothesis that these arise from a partly shared substrate (in particular medial parietal regions), but with the LT bias additionally being mediated by a more extended network, including early sensory and frontal cortices as suggested previously. Consolidating the previous studies is complicated by the different designs and exposure durations used in the previous work. To overcome this problem we performed this comparison within participants using the same stimuli and design for both paradigms. Following our previous work (Park and Kayser 2019), we combined a multisensory ventriloquism task with temporally precise neuroimaging (EEG) and applied single-trial neuro-behavioral modelling to determine the neurophysiological correlates of the ST and LT ventriloquism aftereffect biases.

## METHODS

### Participants

20 right-handed healthy young adults (age range: 22 – 39, mean ± SD: 26.7 ± 4.20, 10 females) participated in the study. All reported normal vision and hearing, with no history of neurological or psychiatric disorders. Each provided written informed consent and were compensated for their time. The study was approved by the local ethics committee of Bielefeld University. One participant’s data was excluded due to not being able to follow the task instructions. Therefore we report data from 19 participants.

### Stimuli

The acoustic stimulus was a 1300 Hz sine wave tone (50 ms duration) sampled at 48 kHz and presented at 64 dB r.m.s. through one of 5 speakers (MKS-26/SW, MONACOR International GmbH & Co. KG, Bremen, Germany) which were located at 5 horizontal locations (−23.2°, – 11.6°, 0°, 11.6°, 23.2°, vertical midline = 0°; negative = left; positive = right). Sound presentation was controlled via a multi-channel soundcard (Creative Sound Blaster Z) and amplified via an audio amplifier (t.amp E4-130, Thomann Germany). Visual stimuli were projected (Acer Predator Z650, Acer Inc., New Taipei City, Taiwan) onto an acoustically transparent screen (Screen International Modigliani, 2×1 m), which was located at 135 cm in front of the participant. The visual stimulus was a cloud of white dots distributed according to a two dimensional Gaussian distribution (N = 200 dots, SD of vertical and horizontal spread 2°, width of a single dot = 0.12°, duration = 50 ms). Stimulus presentation was controlled using the Psychophysics toolbox (Brainard 1997) for MATLAB (The MathWorks Inc., Natick, MA) with ensured temporal synchronization of auditory and visual stimuli.

### Paradigm and task

The paradigm was based on a single-trial audio-visual localization task (Park and Kayser 2019; Wozny and Shams 2011), with trials and conditions designed to probe both the ventriloquism effect and the ventriloquism aftereffect. Participants were seated in front of an acoustically transparent screen with their heads on a chin rest. They were instructed not to move their head while performing the task. Five speakers were located immediately behind the screen and participants responded with a mouse. Their task was to localize a sound during either Audio-Visual (AV: sound and visual stimulus presented simultaneously) or Auditory (A: only sound), trials, or to localize a visual stimulus during Visual trials (V: only visual stimulus). For the AV trials, the locations of auditory and visual stimuli were drawn semi-independently from the 5 locations to yield 6 different audio-visual discrepancies (abbreviated ΔVA; −34.8°, −23.2°, −11.6°, 11.6°, 23.2°, 34.8°). For the A or V trials, stimulus locations were drawn from the 5 locations randomly.

### Experimental setup

Each participant underwent two sessions on different days, in pseudo-randomized order: one for long-term (LT) and one for short-term recalibration (ST). The LT paradigm comprised two parts, 3 consecutive left-wards recalibration blocks, in which the audio-visual discrepancy was always negative (ΔVA < 0°: −34.8°, −23.2°, −11.6°), and 3 consecutive right-wards recalibration blocks in which the discrepancy was always positive (ΔVA > 0°: 11.6°, 23.2°, 34.8°). Other than the negative/positive constraint the positions of the acoustic and visual stimuli were chosen randomly. The ST paradigm comprised 5 blocks, with each block featuring all six audiovisual discrepancies in random sequence. Each audio-visual discrepancy (ΔVA) was repeated 72 / 60 times respectively (LT / ST), resulting in a total of 864 (720) AV and A trials for LT (ST). Around 7% of trials (72 for LT, 55 for ST) were visual-only, interleaved to maintain attention (V trials always came after A trials, thus not interrupting the AV-A sequence). The order of trials was pseudo-randomized. Each trial started with a fixation period (uniform 1100 ms – 1500 ms), followed by the stimulus (50 ms). After a random post-stimulus period (uniform 600 ms – 800 ms) a horizontal bar was shown, along which participants could move a cursor (Figure 1A). A letter indicated which stimulus participants had to localize. On the A trials, participants also reported their confidence by moving a vertical bar between 0% – 100%. There were no constraints on response times, however participants were instructed to respond intuitively, and to not dwell on their response. Inter-trial intervals varied randomly (uniform 800 ms – 1200 ms). Participants were asked to maintain fixation during the entire trial except the response, during which they could freely move their eyes. Eye-tracking data was acquired with a head-mounted eyetracker (EyeLink II, SR Research) at a frequency of 200 Hz. Saccadic eye movements were detected using the ‘cognitive’ setting in the EyeLink II software.

**Figure 1.**
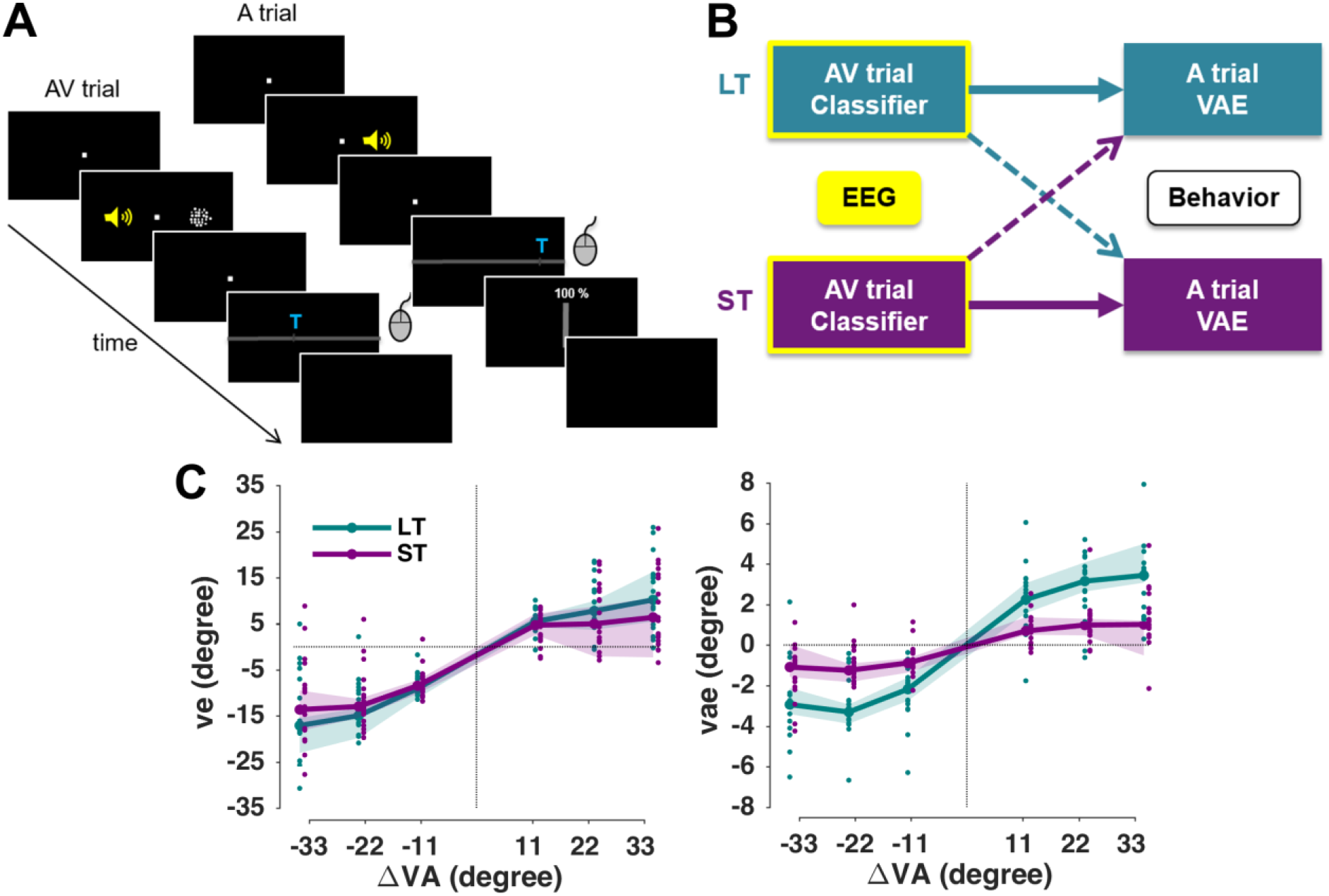
Experiment setup and analyses. (A) Example sequence of AV and A trials (rare V trials are not shown). The yellow speaker is for illustration only; the sound came from speakers placed behind the screen. The participant submitted their response by moving a mouse cursor to the location where they perceived the sound. The confidence rating was only taken in the A trial. For a detailed timing of a sequence, see the main text. (B) In two separate analyses we quantified the predictive power of EEG derived representations of either the multisensory discrepancy or the response in the AV trial to predict the trial-wise *vae* bias in the A trial, either i) within a paradigm (thick arrows) or ii) across-paradigms (dotted arrows). (C) Behavioral results. (left) ventriloquism effect (right) ventriloquism aftereffect, both median across participants (n = 19), shaded areas are 95% confidence intervals around median. Dots show individual participant’s data.

### Analyses of behavioral data

The trial-wise ventriloquism effect (*ve*) in the AV trials was defined as the difference between the actual sound location (A_AV_) and the reported location (R_AV_): *ve* = R_AV_ – A_AV_. The trial-wise ventriloquism aftereffect (*vae*) in the A trials was defined as the difference between the reported location (R_A_) and the mean reported location for all A trials of the same stimulus position (μR_A_), i.e., (*vae* = R_A_ – μR_A_). This ensured that any intrinsic general bias in sound localization would not influence this measure (Wozny and Shams 2011). To quantify the dependency of individual participant’s biases on the audio-visual discrepancy (ΔVA), we fit the trial-wise *ve/vae* biases for each paradigm and participant with the following model:

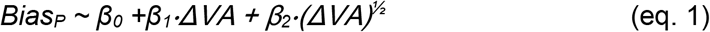

where *Bias* is *ve* or *vae*, and *P* denotes the paradigm, either LT or ST. Here *β_1_, β_2_* quantify the magnitudes of the participant-specific biases. Fitting was done using a maximum likelihood procedure in Matlab R2017a (fitglme.m). Here, in addition to using a linear dependency between bias and discrepancy, we allowed for a non-linear dependency by adding a squareroot term (ΔVA^1/2^). This nonlinear term follows predictions from multisensory causal inference models, which posit that the perceptual bias decreases when the stimuli are sufficiently far apart and don’t seem to originate from a common source (Cao et al. 2019; Körding et al. 2007; Rohe and Noppeney 2015). In a second step, we fit a generalized linear mixed-effects model across all trials from all participants and paradigms. This model extended eq. 1 by adding the paradigm and its interaction with the discrepancy terms and by including participants (*subj*) as random effects to directly compare the group-level biases between LT and ST paradigms:

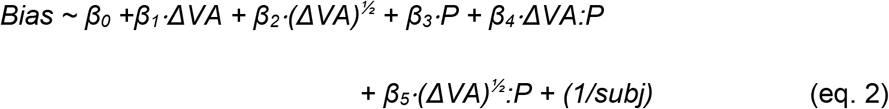

where *Bias* can be *ve* or *vae, P* is the paradigms (LT or ST, coded as categories). The coefficients *β_1_, β_2_* quantify the group-level biases, and *β_4_, β_5_* their interactions with the paradigm.

As previous work has shown that the preceding response can potentially be a driving factor for the ventriloquism aftereffect (Park, Nannt, and Kayser 2020), and because serial-dependencies in perceptual choices prevail in many laboratory paradigms (Fritsche, Mostert, and de Lange 2017; Kiyonaga et al. 2017; Talluri et al. 2018), we tested whether including the previous response would improve the predictive power of model 2:

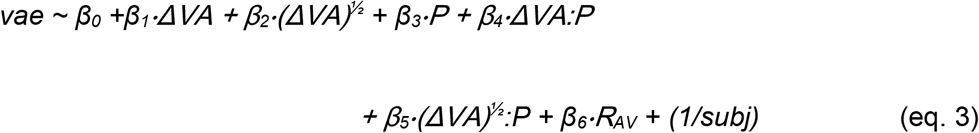

We compared the two models (eq. 2, eq. 3) based on their respective BIC’s.

### EEG acquisition & preprocessing

EEG data were recorded using an active 128 channel Biosemi system (BioSemi, B. V., The Netherlands), with additional four electrodes placed near the outer canthi and below the eyes to record the electro-oculogram (EOG). Electrode offset was below 25 mV. Offline preprocessing and analyses were performed with MATLAB R2017a (The MathWorks, Natick, MA, USA) using the Fieldtrip toolbox (ver. 20190905) (Oostenveld et al. 2011). The data were band-pass filtered between 0.6 and 90 Hz, resampled to 150 Hz and epoched from −0.8 s ∼ 0.65 s around stimulus onset time. Noise removal was performed using ICA simultaneously across all blocks recorded on the same day. The ICA was computed based on 40 PCA components. We removed ICA components that reflect eye movement artefacts, localized muscle activity or poor electrode contacts (17.2 ± 4.45 rejected components per participant, mean ± SD). These were identified as in our previous studies (Grabot and Kayser 2020; Kayser, Philiastides, and Kayser 2017) following definitions provided in the literature (Hipp and Siegel 2013; O’Beirne and Patuzzi 1999).

### EEG discriminant analysis

To extract neural signatures of the encoding of different variables of interest we applied a cross-validated regularized linear discriminant analysis (LDA) (Blankertz et al. 2011; Parra and Sajda 2003) to the single trial data from the AV trials. Preprocessed EEG data were filtered between 2 Hz and 40 Hz (4th order Butterworth filter) and the LDA was applied to the data aligned to stimulus onset (0 s) in 40 ms sliding windows, with 6.7 ms time-steps (time window: −0.4 s ∼ 0.5 s). The regularization parameter was set to 0.1 as in previous work (Park and Kayser 2019).

We computed separate linear discriminant classifiers for the two variables of interest: i) ΔVA, and ii) the response in the AV trial (R_AV_). For each variable we classified whether that variable was left- or right-lateralized by grouping the single trial values into left (< 0°) or right (> 0°), similar to our previous study on the ventriloquism aftereffect (Park and Kayser 2019). Importantly, by binarizing the variables in this way, we also avoided specific assumptions about whether the aftereffect follow allows a linear or non-linear dependency on AV discrepancy. The classifier performance was characterized as the ROC (Receiver operating characteristic)’s AUC (area under curve) obtained from 6-fold cross-validation. We derived scalp topographies for each classifier by estimating the corresponding forward model, defined as the normalized correlation between the discriminant component and the EEG activity (Kayser et al. 2017; Parra et al. 2005).

### Neuro-behavioral models predicting the trial-wise aftereffect

We then used these classifiers in neuro-behavioral models to elucidate the correlates of the single trial *vae* biases. We implemented two analyses that differed in the trials used to train the classifier and the trials used to predict the behavioral bias: first, we tested the ability to predict the *vae* bias in the A trial based on the EEG activity obtained in the preceding AV trial within each paradigm (Figure 1B, thick arrows); second, we tested the ability to cross-predict the *vae* bias in one paradigm (e.g. ST) based on the brain activity in the AV trials of the other paradigm (eg. LT; Figure 1B, dotted arrows). These two analyses were geared to reveal the cerebral representations of audio-visual disparity (or response behavior) in the multisensory trial that are predictive of the response bias in the subsequent unisensory trial, specifically within a paradigm or consistently across paradigms. The cross-classification analysis directly tests the assumption that the cerebral activations (here captured by the classifier weights) representing the audio-visual discrepancy and driving the aftereffect are identical across paradigms both in their spatial generators within the brain and in time relative to the presentation of the AV stimulus.

We computed two linear models for each of the two analyses (within or between paradigms), with LDA-ΔVA (or LDA-R_AV_) standing for the respective continuous single-trial classifier predictions, which provides a proxy to the cerebral representation of the respective variable of interest (Kayser and Kayser 2020; Kayser, McNair, and Kayser 2016; Park and Kayser 2019; Philiastides, Heekeren, and Sajda 2014):

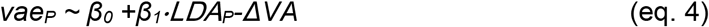

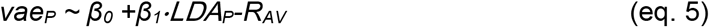

where *P* denotes paradigms (LT, ST). From the coefficients (*β_1_*) obtained for individual participants we then determined i) whether the cerebral encoding of a variable offered significant predictive information for the *vae* by testing the coefficient at the group-level against zero, ii) when this prediction emerged, and iii) by looking at the forward models of the respective LDAs, we determined the underlying cerebral sources.

These models were computed using EEG activity from the stimulus- and post-stimulus period in the AV trial based on 3-fold cross-validation, using distinct folds of trials to determine the weights of the LDA and to compute the regression models by applying these weights to activity in a different fold (or paradigm). We averaged the resulting betas across 30 repeats of this analysis. We then computed group-level t-values for the coefficients for each predictor at each time point, and assessed their significance using cluster-based permutation statistics controlling for multiple comparisons (below: Statistical analysis).

### EEG source analysis

Single-trial source signals were derived using a linear constrained minimum variance beamformer (LCMV, 7% normalization, using a covariance matrix obtained from −0.6 s ∼ 0.5 s peri-stimulus period, projecting along the dominant dipole orientation) as implemented in the FieldTrip toolbox (Oostenveld et al. 2011). As participant-specific anatomical data were not available, we used a standardized head model using the average template brain of the Montreal Neurological Institute. Lead fields were computed using a grid spacing of 6 mm. Then, we computed the source-level correlation between the single-trial grid-wise source activity for each participant and the LDA output activity over trials in order to quantify the relevant source regions at specific time points, similar to obtaining the forward scalp distributions by correlating the sensor and LDA components (Haufe et al. 2014; Parra et al. 2005). Source correlations were z-scored before averaging across participants. To interpret these group-level source maps we thresholded these above the 95th percentile, and identified clusters with a minimum cluster size of 80 voxels based on a connected components algorithm (SPM8 toolbox, 2008 Wellcome Trust Centre for Neuroimaging). We then extracted the anatomical labels based on the AAL atlas (Tzourio-Mazoyer et al. 2002), to determine those regions covered by these clusters (reporting atlas regions containing at least 20 voxels and occupying at least 30% of the total number of voxels for each atlas region).

### Eye movement analyses

We checked for excess eye-movements by computing the number of saccades between −50 ms ∼ 100 ms of stimulus onset that were larger than 1 deg visual angle. We also computed the percentage of saccades in AV trials between stimulus offset and 400 ms that pointed in the same direction as ΔVA. Finally, to rule out the possibility that eye movements contribute by inducing specific artifacts in the EEG signals, we applied the neuro-behavioral analyses to the EOG data rather than the EEG data.

### Statistical analysis

To test the (trial-averaged) *ve* and *vae* from zero we used a sign rank test, correcting for multiple tests using the Holm procedure with a family-wise error rate of p < 0.05 (Figure 1C). The confidence intervals for the median (e.g. Figure 1C, Figure S2A) were obtained using the bootstrap hybrid method with 199 resamples (Bootstrap Matlab Toolbox, Zoubir and Boashash 1998). Group-level inference on the LDA time-course was performed using randomization procedures and cluster-based statistical enhancement controlling for multiple comparisons along time (Maris and Oostenveld 2007; Nichols and Holmes 2002). First, we shuffled the sign of the true single-participant effects (the signs of the chance-level corrected AUC values; or the signs of single-participant regression betas) and obtained distributions of group-level effects (mean for AUC, t-values for regression models) based on 3000 randomizations. We then applied spatial clustering based on a minimal cluster size of 4 and using the sum as cluster-statistics. For testing the LDA performance, we thresholded the first-level effects based on the 99th percentile (i.e. p < 0.01) of the full distribution of randomized AUC values. For testing regression betas, we used parametric thresholds corresponding to a two-sided p < 0.01 (tcrit = 2.81, d.f. = 18). The threshold for determining significant clusters was p < 0.01 (two-sided). We tested for significant temporal clusters for classifier performance in the whole time window of interest (−0.4 s ∼ 0.5 s), while the neuro-behavioral models were restricted to a time window of interest within the significant discriminant performance for the respective variable (ΔVA, R_AV_). For the cross paradigm analysis (Figure 2C), we computed a conjunction statistics, obtained at each time point by taking the smaller of the two t-values obtained from LDA_ST_ → *vae*_LT_ and LDA_LT_ → *vae*_ST_ (eq. 4, 5) (Nichols et al. 2005).

**Figure 2.**
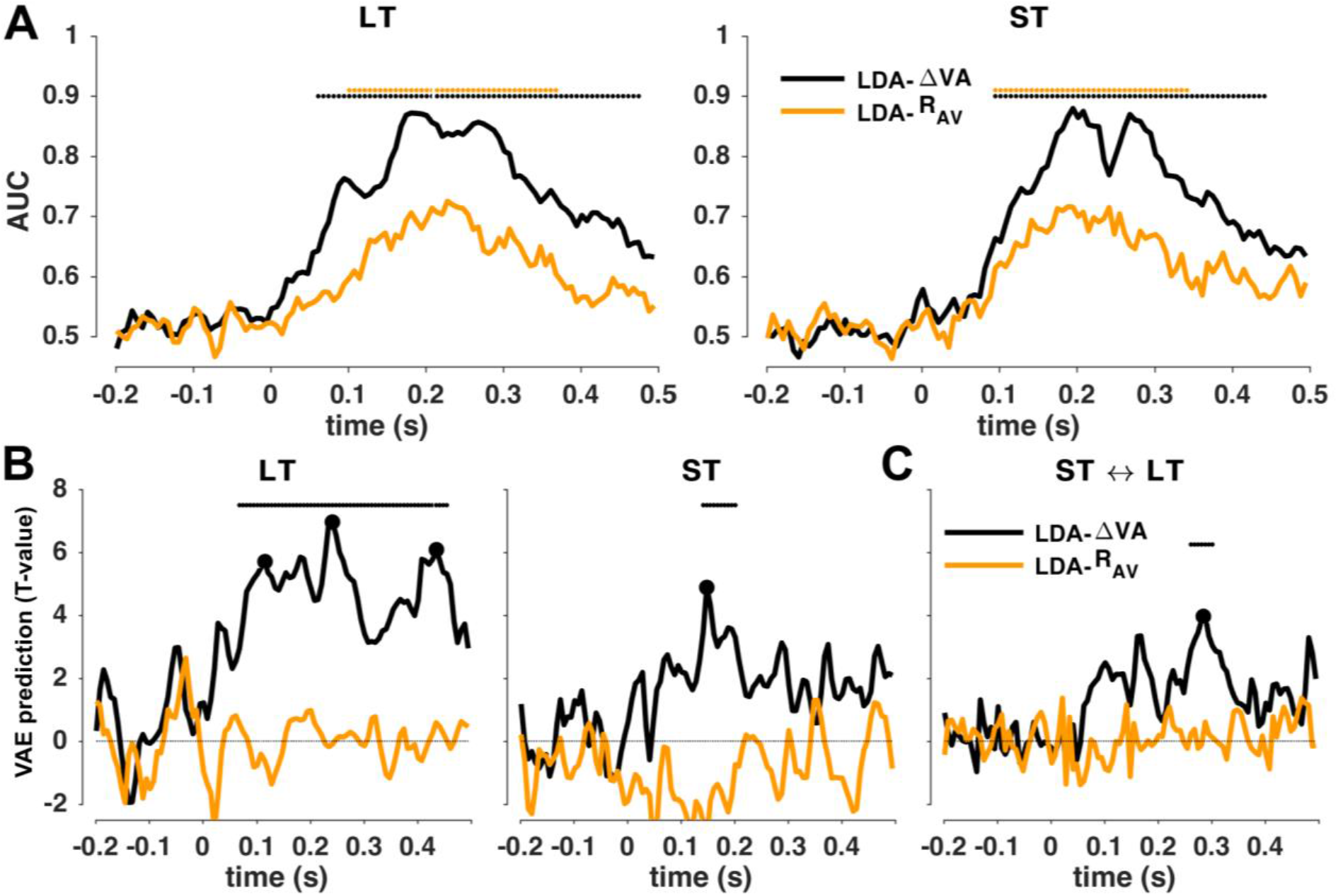
Predicting the trial-wise aftereffects based on neurophysiological representations. (A) Classifier performance (group-level mean, n = 19) for both paradigms (short-term ST; long-term LT) as cross-validated area under the ROC curve (AUC). (B) Neuro-behavioral models predicting the trial-wise aftereffect within paradigms based on the EEG-derived cerebral encoding of sensory (ΔVA) or motor (R_AV_) variables in the AV trial. Graphs show group-level t-maps of the underlying regression betas. (C) Neuro-behavioral models predicting the aftereffect across paradigms. Significance based on clusterbased permutation-based statistics (p < 0.01; see Methods).

To compare the similarities of the group-level forward models of the LDA classifiers obtained in different paradigms, or at different time points, we quantified their group-level similarity using Pearson correlation. Statistical significance was tested using bootstrapping over the (random) selection of participants used to compute the group-level mean (at p < 0.01, using 3000 resamples).

## RESULTS

### Behavioral biases

Behavioral responses in AV trials revealed a clear ventriloquism bias as a function of the audiovisual discrepancy (ΔVA = V_AV_ – A_AV_), reflecting the influence of the visual stimulus on the perceived location of the simultaneous sound (Figure 1C, left). All group-level *ve* biases were significantly different from zero (sign rank test: p < 0.01 for all 6 ΔVA). A GLMM revealed that the ventriloquism bias varies nonlinearly with the discrepancy but does not differ between paradigms (Table 1A, see also Figure S1A and Table S1 for single participant effects).

**Table 1:** Generalized linear mixed models (eq. 2, 3) results for fixed effects. CI: 95% confidence interval (parametric); BIC: Bayesian information criterion; AIC: Akaike information criterion; LL: Loglikelihood. (A) Reveals the linear and non-linear dependency of the *ve* on multisensory discrepancy (ΔVA), which did not differ between paradigms. (B) Reveals the linear and non-linear dependency of the *vae* on multisensory discrepancy, which both differed between paradigms. (C) Comparing models 2 and 3 shows that some of the variance in the aftereffect is also explained by the response in the AV trial (R_AV_).

| **(A) Eq. 2: $v_e \sim \beta_0 + \beta_1 \Delta VA + \beta_2 (\Delta VA)^{1/2} + \beta_3 P + \beta_4 \Delta VA:P + \beta_5 (\Delta VA)^{1/2}:P + (1/subj)$** |  |  |  |  |  |  |
| --- | --- | --- | --- | --- | --- | --- |
| Name | Estimate ($\beta$) | t-statistics | p-value | CI (95%) | | Model fits |
| Intercept | -2.0665 | -3.1630 | 0.0016 | -3.3471 | -0.7859 | BIC: |
| $(\Delta VA)^{1/2}$ | 2.2056 | 7.7459 | 0.0000 | 1.6474 | 2.7637 | 109000 |
| $\Delta VA$ | 0.0744 | 1.3598 | 0.1739 | -0.0329 | 0.1817 | AIC: |
| $P$ | 0.0636 | 0.4070 | 0.6840 | -0.2427 | 0.3700 | 108940 |
| $(\Delta VA)^{1/2}:P$ | -0.1427 | -0.7676 | 0.4427 | -0.5070 | 0.2216 | LL: |
| $\Delta VA:P$ | -0.0513 | -1.4370 | 0.1507 | -0.1214 | 0.0187 | -54460 |
| **(B) Eq. 2: $v_{ae} \sim \beta_0 + \beta_1 \Delta VA + \beta_2 (\Delta VA)^{1/2} + \beta_3 P + \beta_4 \Delta VA:P + \beta_5 (\Delta VA)^{1/2}:P + (1/subj)$** |  |  |  |  |  |  |
| Intercept | 0.0023 | 0.0134 | 0.9893 | -0.3297 | 0.3342 | BIC: |
| $(\Delta VA)^{1/2}$ | 1.6087 | 7.9828 | 0.0000 | 1.2137 | 2.0037 | 98656 |
| $\Delta VA$ | -0.1368 | -3.5306 | 0.0004 | -0.2127 | -0.0608 | AIC: |
| $P$ | -0.0006 | -0.0053 | 0.9958 | -0.2172 | 0.2161 | 98595 |
| $(\Delta VA)^{1/2}:P$ | -0.7178 | -5.4568 | 0.0000 | -0.9757 | -0.4600 | LL: |
| $\Delta VA:P$ | 0.0733 | 2.8981 | 0.0038 | 0.0237 | 0.1229 | -49290 |
| **(C) Eq. 3: $v_{ae} \sim \beta_0 + \beta_1 \Delta VA + \beta_2 (\Delta VA)^{1/2} + \beta_3 P + \beta_4 \Delta VA:P + \beta_5 (\Delta VA)^{1/2}:P + \beta_6 R_{AV} + (1/subj)$** |  |  |  |  |  |  |
| Intercept | 0.0542 | 0.3198 | 0.7491 | -0.2778 | 0.3862 | BIC: |
| $(\Delta VA)^{1/2}$ | 1.5506 | 7.6945 | 0.0000 | 1.1556 | 1.9457 | 98631 |
| $\Delta VA$ | -0.1256 | -3.2412 | 0.0012 | -0.2015 | -0.0496 | AIC: |
| $R_{AV}$ | 0.0262 | 5.8954 | 0.0000 | 0.0175 | 0.0349 | 98562 |
| $P$ | -0.0002 | -0.0015 | 0.9988 | -0.2166 | 0.2162 | LL: |
| $(\Delta VA)^{1/2}:P$ | -0.7137 | -5.4320 | 0.0000 | -0.9713 | -0.4562 | -49272 |
| $\Delta VA:P$ | 0.0746 | 2.9523 | 0.0032 | 0.0251 | 0.1241 | |

Regarding the ventriloquism aftereffect, the behavioral responses in the A trials revealed a clear bias in the direction of the previous trial’s ΔVA (Figure 1C, right). All group-level *vae* biases were significantly different from zero (sign rank test: p < 0.01 for all 6 ΔVA). The GLMM showed that the aftereffect exhibits both a linear and a nonlinear dependence on discrepancy (Table 1B, cf. Figure S1B and Table S1 for single participant effects). Importantly, both the linear and nonlinear dependency on ΔVA differed between paradigms (p < 0.01; Table 1B).

Starting from the notion that the aftereffect is a direct cause of the ventriloquism effect, we probed the trial-by-trial link of these biases by quantifying their within-participant correlation. The median Spearman’s rank correlation coefficients for LT and ST were 0.382 and 0.211 respectively. A permutation test on the difference of medians confirmed that the correlation was stronger in the long-term paradigm LT (p = 0.00016, based on a 50000 bootstrap samples, Figure S2A). Closer inspection of the trial-wise dynamics of these effects revealed a clear accumulation of the aftereffect over the course of the long-term but not over the short-term paradigm (Figure. S2B).The ventriloquist bias in the AV trials, in contrast, did not change in strength over time (Figure. S2B). These results show that prolonged exposure to a consistent audio-visual discrepancy results in a stronger bias, possibly resulting from a stronger trial-wise link between the ventriloquism bias in the AV trial and it’s persistent influence on the subsequent sound.

### The aftereffect bias reflects the previous multisensory discrepancy

Previous studies suggested two potential factors driving the aftereffect: the sensory discrepancy (ΔVA) in the previous trial, or the participant’s response in that trial (R_AV_) (Park and Kayser 2019; Van der Burg, Alais, and Cass 2018). Indeed, in many laboratory paradigms sequential effects between the responses on different trials emerge, by which the previous response is predictive of the subsequent one (Fritsche et al. 2017; Kiyonaga et al. 2017; Talluri et al. 2018; Urai et al. 2019). We asked whether the single-trial aftereffect biases are better accounted for by allowing a dependency on the previous response R_AV_ (eq. 2 vs. eq. 3; Table 1B, C). The model fit improved by adding the previous response (ΔBIC = 25), suggesting that the previous response contributes to shaping the trial-wise bias in the AV trial in addition to the multisensory discrepancy. For the following analysis of the neurophysiological correlates we considered both ΔVA and R_AV_ as variables of interest whose cerebral representations in the AV trial could be predictive of the subsequent aftereffect.

### Predicting the aftereffect from neurophysiological representations

To probe whether and which EEG activations reflecting the cerebral encoding of task-relevant variables (e.g. ΔVA) in the AV trial are predictive of the subsequent *vae* bias, we extracted EEG-derived representations of these variables using single-trial classification. We then quantified whether and which of these representations are predictive of the single trial *vae* bias. In a first analysis, we tested this within the LT and ST paradigms individually, to potentially reveal representations that are either paradigm-specific or possibly exhibit common properties (time, topographies) between paradigms. In a second analysis, we directly aimed to extract EEG-derived representations that are common to both paradigms, by predicting the bias in one paradigm based on classifiers trained on the EEG activity in the other paradigm.

We applied linear discriminant analysis (LDA) to the AV trial data to probe when the EEG activity allows the (cross-validated) classification of the two main variables of interest: ΔVA and R_AV_. Here, ΔVA served as the main variable of interest driving the aftereffect, and R_AV_ as a control. In both the LT and ST paradigms, discrimination performance became significant from around 100 ms post stimulus onset (Figure 2A). The performances of both classifiers were significant over a long time in the LT (LDA-ΔVA: p = 0.0003, t_cluster_ = 16.71, peak = 0.87, range = [62 ms, 475 ms]; LDA-R_AV_: p = 0.0003, t_cluster_ = 6.70, peak = 0.73, range = [102 ms, 368 ms]), and ST paradigm (LDA-ΔVA: p = 0.0003, t_cluster_ = 14.18, peak = 0.88, range = [95 ms, 442 ms]; LDA-R_AV_: p = 0.0003, t_cluster_ = 6.35, peak = 0.72, range = [95 ms, 342 ms]).

We then asked whether and when the cerebral representations of these variables are predictive of the aftereffect bias. Figure 2B shows the respective group-level t-values of the regression betas from eq. 4, 5 for the within-paradigm analysis (see Figure S3 for time courses of the mean rather than significance). The LDA-ΔVA predicted the subsequent trial-wise *vae* bias between 75 ms ∼ 475 ms in the LT paradigm (p = 0.0003, t_cluster_ = 311.6, t_peak_ = 6.96, Cohen’s *d* = 1.60). In the ST paradigm, the LDA-ΔVA predicted the bias between 142 ms ∼ 202 ms (p = 0.001, t_cluster_ = 36.1, t_peak_ = 4.88, Cohen’s *d* = 1.12; Figure 2B, left), with the significant clusters overlapping between both paradigms. In contrast, the LDA-R_AV_ in the AV trial was not predictive of the bias in either paradigm (no significant clusters; maximum Cohen’s *d* = 0.22, at 202 ms in the LT, d = 0.30 at 355 ms for the ST).

Then, in a direct cross-decoding analysis we tested whether cerebral representations of these variables can predict the bias between paradigms (Figure. 2C). This revealed a significant cluster between 261 ms and 301 ms, in which the LDA-ΔVA in the AV trial of the LT paradigm predicts the bias in the A trial in the ST paradigm, and vice versa (obtained from the conjunction statistics cross-predicting in both directions; p = 0.0003, t_cluster_ = 23.8, t_peak_ = 3.99, Cohen’s d = 0.91).

### Distinct neurophysiological sources of the short- and long-term biases

To better understand the physiological correlates of the aftereffect biases, we extracted key time points of interest, defined as the local and global peaks in the neuro-behavioral analysis (Figures 2B, C black dots). We then investigated the underlying neural generators by inspecting the LDA forward models and source maps. From the within-LT analysis we derived three time points (local peaks at 115 ms and 435 ms; global peak at 241 ms). These time points were specific to the LT paradigm, as the respective EEG activity in the ST paradigm at these moments was not predictive of the aftereffect (at an uncorrected p < 0.01: t = 1.85, 1.87, 1.69 and Cohen’s *d*: 0.4237, 0.4299, 0.3870). From the within-ST analysis we derived one time point (148 ms) (the LT analysis revealed a significant cluster at the same time). From the cross-paradigm analysis we obtained one time point (global peak at 288 ms). Given that the significant clusters for the within ST and LT analysis overlapped, we asked whether the forward models of the LDA-ΔVA components were similar (at 148 ms): these were indeed highly correlated (Spearman’s ρ = 0.97, Bootstrap-based CI = [0.52 0.98], p < 0.001), suggesting that the underlying generators are similar. We hence combined the topographies and sources across paradigms at 148 ms and 288 ms. The resulting forward topographies are shown in Figure 3A, B.

**Figure 3.**
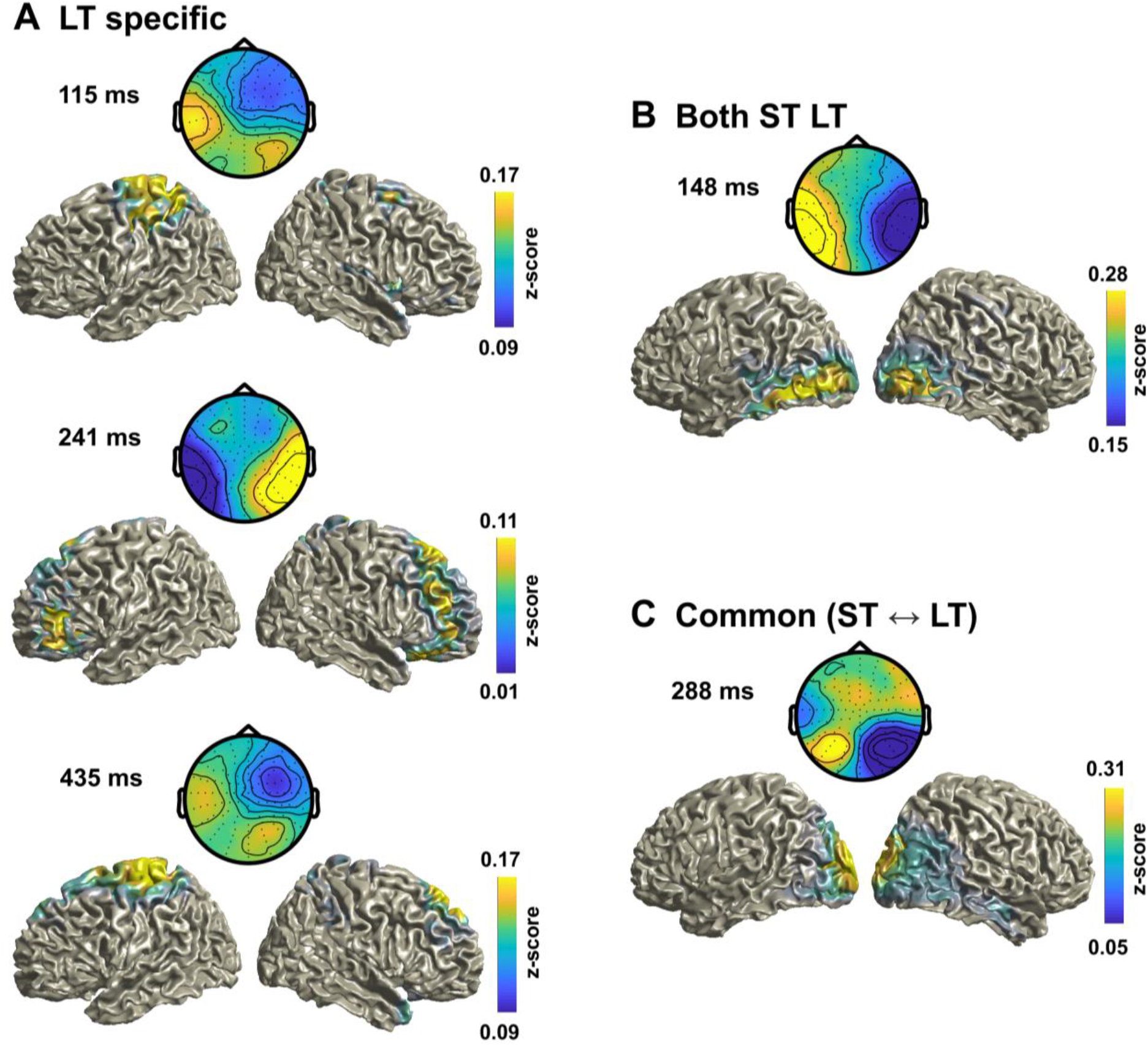
EEG topographies and source maps for the LDA-ΔVA classifier. Group-averaged topographies (forward models) and source maps for (A) the three LT specific time points derived in Figure 2B. (B) Time point common to both paradigms, and (C) for the peak time point in the cross-paradigm analysis in Figure 2C. The data are shown as z-score transformed correlation from the source LDA activity and single trial response data (see Methods). For B) and C) the data were averaged across paradigms.

Then, we asked whether the relevant neurophysiological sources were similar between time points within paradigms (LT: 115 ms vs. 241 ms: Rho = −0.34, CI = [−0.79, 0.29]; 115 ms vs. 435 ms: Rho = 0.86, CI = [−0.42, 0.97]; 241 ms vs. 435 ms: Rho = −0.43, CI = [−0.81, 0.59]) ST/LT: 148 ms vs. 288 ms: Rho = 0.50, CI = [−0.45, 0.75]). The group-level forward models were not significantly correlated between time points (all pairs p > 0.05, group-level bootstrap confidence intervals). This demonstrates that activity at each time point reflects distinct neurophysiological contributions to the aftereffect, suggesting a contribution from multiple and temporally dispersed processes. Furthermore, this result demonstrates that partly distinct processes contribute to the short- and long-term biases.

Finally, we inspected the underlying generators in source space. The group-level source maps revealed an involvement of medial superior parietal regions (in particular the precuneus) at multiple time points and common to both paradigms (e.g. at 288 ms; Figure 3B,C), in line with the hypothesis that parietal structures involved in sensory causal inference and memory mediate recalibration in general. Common to both paradigms were also sources in sensory regions (occipital and temporal cortex; at 148 ms and 288 ms), while sources specific to the LT paradigm involved precentral and frontal regions (Figure 3A, Table 2).

**Table 2.** Anatomical labels of source clusters for each time point in Figure 3. Anatomical labels are based on the AAL atlas (Tzourio-Mazoyer et al. 2002). See Methods for the extraction of these regions.

| 115 ms | 148 ms | 241 ms | 288 ms | 435 ms |
| --- | --- | --- | --- | --- |
| LT | LT/ST | LT | ST ↔ LT | LT |
| Pre-/Para-/ | Temporal Inf | Frontal Inf/Mid/Sup | Temporal Inf/Mid | Frontal Sup |
| Post-central | Occipital Inf/Mid | Supp. motor | Cuneus | Supp. Motor |
| Precuneus | Calcarine | Paracentral | Precuneus | Pre-/Para-/ |
| Parietal Inf/Sup |  | Parietal Sup | Angular | Post-central |
|  |  |  | Calcarine | Parietal Inf/Sup |
|  |  |  | Occ. Inf/Mid/Sup |  |

### Eye movements do not confound these results

To ensure that potential eye-movements do not confound any of these conclusions, we analyzed both eye movements themselves and their predictive power for the single trial biases (Kopco et al. 2009; Werner-Reiss et al. 2003). First, only a few A trials (2.3% ± 0.5% mean ± SEM across participants and trials) contained saccadic eye movements during stimulus presentation (> 1 deg; between −50 ms ∼ 100 ms of stimulus onset), showing that participants maintained fixation well. Second, we computed the percentage of saccades in AV trials between stimulus onset and 400 ms that pointed in the same direction as ΔVA, and hence would directly confound with the direction of ΔVA as a predictor. Overall, the direction of saccades was very balanced: with only 51.2% ± 3.1% (mean ± SEM. LT: 52.0% ± 3.1%, ST: 50.3 ± 3.2%) pointing in the same direction as ΔVA. Finally, to rule out the possibility that eye movements contribute by inducing specific artifacts to the EEG analysis, we implemented the neuro-behavioral analyses using the EOG data. This did not provide any significant relation between EOG activity and the *vae* bias (using the same statistical criteria as for the EEG data; Figure S4).

## DISCUSSION

We often encounter seemingly discrepant sensory information, such as when watching a movie on a screen and hearing sounds through earphones. Our brain reconciles such discrepant sensory information by adapting to this sensory disparity by recalibration over distinct time scales. Despite being a well-studied behavioral phenomenon, the neural underpinnings of multisensory spatial recalibration remain debated, in particular whether the behavioral biases from long- and short-term exposure to discrepant information arise from largely the same or distinct neural origins. Our results are in line with the hypothesis that the ventriloquism aftereffects of short and long-term exposure are mediated by common neurophysiological correlates that likely arise from sensory and parietal regions involved in multisensory inference and memory. However, they also show that prolonged exposure to consistent multisensory discrepancy additionally recruits frontal regions that reinforce this behavioral effect and which may be responsible for the increased aftereffect bias following long-term exposure.

### The neural underpinnings of the spatial ventriloquism aftereffect

Sensory recalibration as in the ventriloquism aftereffect is robustly seen in a wide range of paradigms after exposure to single stimulus as well as after long-term exposure (Bruns & Röder, 2015, 2019; Lewald, 2002; Mendonça et al., 2015; Recanzone, 1998; Wozny & Shams, 2011). Behavioral studies raised the intriguing hypothesis that the short- and long-term bias may arise from different neural mechanisms, in part as the short-term effect seems to generalize across different attributes of the inducing stimuli (e.g. sound frequencies), while the long-term effect is more specific (Bruns and Röder 2015, 2019; Lewald 2002; Recanzone 1998). However, the degree of stimulus specificity remains debated (Frissen et al. 2003, 2005), and by nature behavioral studies remain inconclusive as to the precise neural mechanisms.

Previous neuroimaging studies reported effects of prolonged exposure to audio-visual discrepancy in sound-evoked EEG responses at around 100 ms after stimulus onset, suggesting a neural correlate near primary auditory cortex (Bruns et al. 2011). While being in line with single neuron data (Recanzone 1998; Recanzone et al. 2000), these studies also postulated that the long-term aftereffect may be mediated by the interplay between auditory cortices and parietal regions involved in multisensory integration (Bruns et al. 2011; Zierul et al. 2017). However, and despite some indirect evidence from studies on functional connectivity (Zierul et al. 2017), this previous work did not demonstrate such a distributed origin of longterm recalibration.

One caveat is that these studies either explicitly focused on auditory cortices or relied on neural signatures of sound encoding (e.g. sound evoked potentials) to probe for a change induced by the exposure to discrepant audio-visual stimuli. By design, this approach likely reveals brain regions primarily involved in sound encoding, while those concerned with other computations, such as multisensory fusion or sensory causal inference, are less likely to emerge. Here we took a different approach, and focused on neural processes reflecting the spatial discrepancy between the auditory and visual stimuli, which, as shown here and previously (Park and Kayser 2019; Wozny and Shams 2011; Zierul et al. 2017), is the key sensory variable driving the aftereffect. Thereby our approach focused on neural processes reflecting the encoding of the sensory cause of the adaptive effect, rather than on neural signatures of the individual stimulus.

Previous work has suggested that adaptive recalibration, in particular on a trial-by-trial level, may not only be related to the previous sensory input but also to variables related to the reported experience, or previous motor response (Park et al. 2020; Van der Burg et al. 2018). Indeed, the two prominent events during the AV trial are the exposure to a discrepant audiovisual stimulus and the corresponding response. Contrasting these two in their predictive power of the subsequent bias, we previously reported in favor of a sensory-rather than motor-related origin of the ventriloquism aftereffect (Park and Kayser 2019; Park et al. 2020; Van der Burg et al. 2018). In the present dataset we found that the previous motor response contributes significantly, in addition to the multisensory discrepancy. Hence we also asked whether neurophysiological signatures of the previous response (R_AV_) could also serve as a predictor of the trial-wise aftereffect. In contrast to the cerebral processes encoding multisensory discrepancy, those reflecting the previous response were not significantly predictive of the aftereffects. This further corroborates the sensory nature of spatial ventriloquism and rules out motor-related confounds as mediators of the reported neurophysiological underpinnings.

### Multiple timescales of the ventriloquism aftereffect

Our results consolidate previous studies on the different time scales of spatial ventriloquism by showing that the trial-by-trial and long-term aftereffects arise largely from shared neural processes, while additional processes are further recruited following long-term exposure. While the sources underlying these EEG data have to be interpreted with care, our results show that the long-term effect is driven by a larger network prominently including parietal and frontal regions. In contrast, occipital sensory and parietal regions mediate the aftereffect consistently following both short- and long-term adaptation. Previously, using MEG-based source-imaging we have shown that the short-term effect is mediated by medial parietal regions (Park and Kayser 2019) involved in spatial working memory (Martinkauppi 2000) and sound localization, such as the precuneus (Lewald et al. 2008; Tao et al. 2015). The present data suggest that these regions mediate the ventriloquism aftereffect over multiple time scales, highlighting that structures involved in spatial working or procedural memory play a central role for this adaptive behavior (Schott et al. 2019) (Mueller 2018). While participants’ task was to localize the sound, we suggest that implicit memory about the previously received discrepant spatial information persists between trials and affects the subsequent perceptual response. This may then involve the previously reported modulation of sound encoding within low-level auditory cortices (Park and Kayser 2019; Zierul et al. 2017).

The long-term bias was also mediated by a more extensive network involving pre-central and frontal regions. Previous work suggested that long-term exposure to discrepant audio-visual stimuli changes connectivity between frontal and auditory cortical regions (Zierul et al. 2017) and has implied a role of inferior frontal regions in multisensory causal inference (Cao et al. 2019; Rohe and Noppeney 2015). In the long-term paradigm the sequence of multisensory discrepancies followed a regular pattern, while it was random in the short-term version. Hence, this systematic pattern may engage regions involved in updating working memory about sensory causal relations, as a consistent discrepancy allows the prospective formation of predictions about upcoming stimuli (Collins and Frank 2013; Curtis 2006; Nee and D’Esposito 2016; Noppeney, Ostwald, and Werner 2010). This regularity of the sensory environment may hence involve frontal regions that exploit this regularity on longer times beyond the more immediate aftereffect arising from parietal regions. Such a divided role of parietal and frontal regions in contributing to multisensory recalibration is in line with the notion that parietal regions contribute to the fusion of multisensory information within a trial, while frontal regions are also engaged in the causal inference of whether different stimuli likely arise from a common source (Cao et al. 2019; Rohe et al. 2019; Rohe and Noppeney 2015).Such an inference process benefits from knowledge about the recent stimulus history (Beierholm et al. 2019). It remains to be understood whether the same or distinct frontal regions contribute to causal inference within a trial and the fostering of recalibration based on the cumulative stimulus history.

Our data also show that the sequential accumulation of bias evident at the level of behavior is specific to the aftereffect, while the ventriloquism bias itself remains stable, even following long-term exposure (Figure 1C, S2B). The ventriloquism aftereffect has been considered a direct consequence of the ventriloquism effect, by which the discrepant audio-visual information is combined into a biased perceived location of the multisensory stimulus (Lewald 2002; Radeau and Bertelson 1977, 1978; Recanzone 1998, 2009; Rohlf et al. 2020). This is corroborated by the significant trial-by-trial correlation of both biases in both paradigms (Figure S2). Still, the stronger trial-wise link in the long-term data suggests that the stronger aftereffect in the long-term paradigm may be a consequence of an enhanced influence from the previous trial with prolonged exposure to consistent audio-visual discrepancies rather than a stronger fusion of such discrepant information in the AV trial itself. This differential dependency on the stimulus history could be used in the future to disentangle the various prefrontal contributions to multisensory integration and recalibration.

## ACKNOWLEDGEMENTS

We thank Bora Kim for helping with collecting the data. This work was supported by the European Research Council (to CK ERC-2014-CoG; grant No 646657).

## Supplementary Information

**Figure S1.**
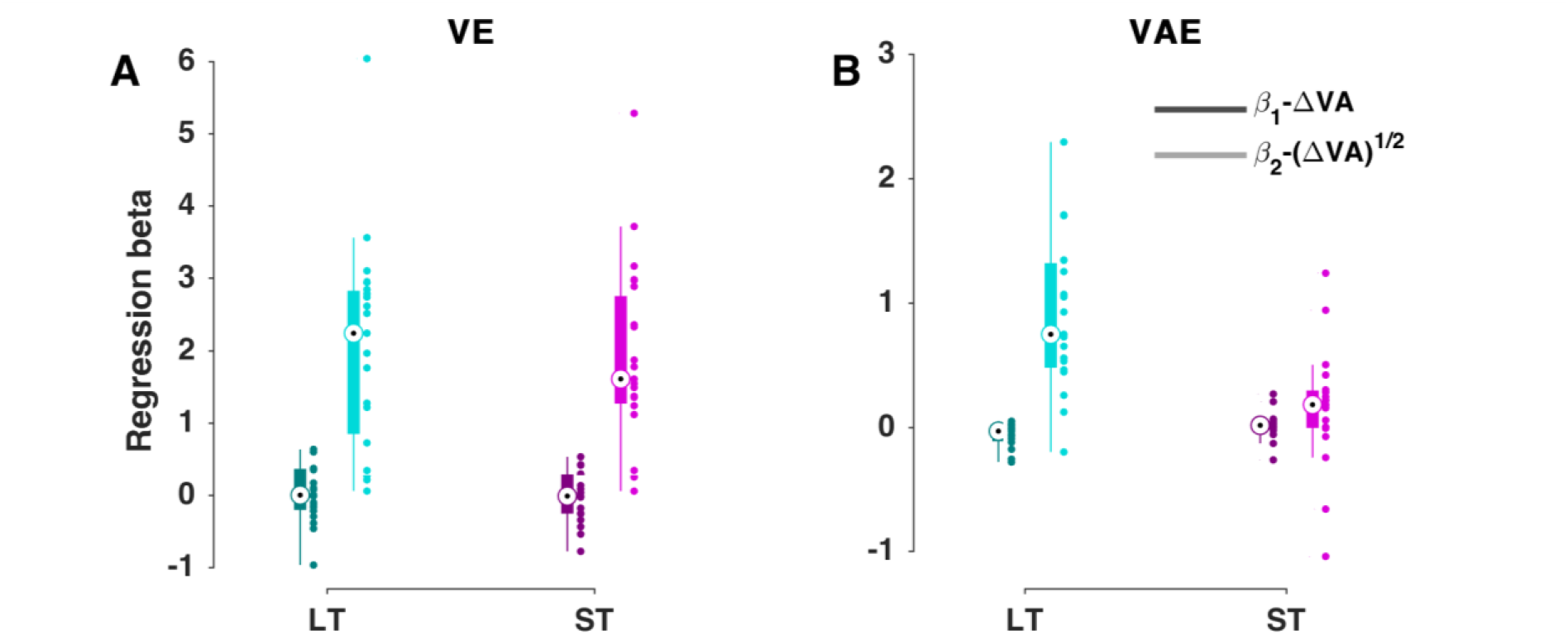
Results from fitting individual participant biases. While in the main text (Table 1) we report the fit of a GLMM across participants and paradigms, we here report results obtained from by fitting individual participants trial-averaged and paradigm-specific biases with a model depending linearly and nonlinearly on the audio-visual discrepancy: *Bias ∼ β_0_ + β_1_ · ΔVA + β_2_·(ΔVA)^½^*. In the graph circles indicate medians, boxes are the 25th and 75th percentiles. Filled dots are individual participants. VE: ventriloquism effect. VAE: ventriloquism aftereffect.

**Table S1.** Medians and interquartile ranges (IQR) for single participant biases from Figure S1. Data are (median, IQR). VE: ventriloquism effect. VAE: ventriloquism aftereffect.

|  |  | VE | VAE |
| --- | --- | --- | --- |
| LT | $(\Delta VA)^{1/2}$ | 2.239, 1.987 | 0.748, 0.837 |
| | $\Delta VA$ | -0.004, 0.569 | -0.032, 0.112 |
| ST | $(\Delta VA)^{1/2}$ | 1.605, 1.491 | -0.018, 0.543 |
| | $\Delta VA$ | 0.181, 0.302 | 0.015, 0.074 |

**Figure S2.**
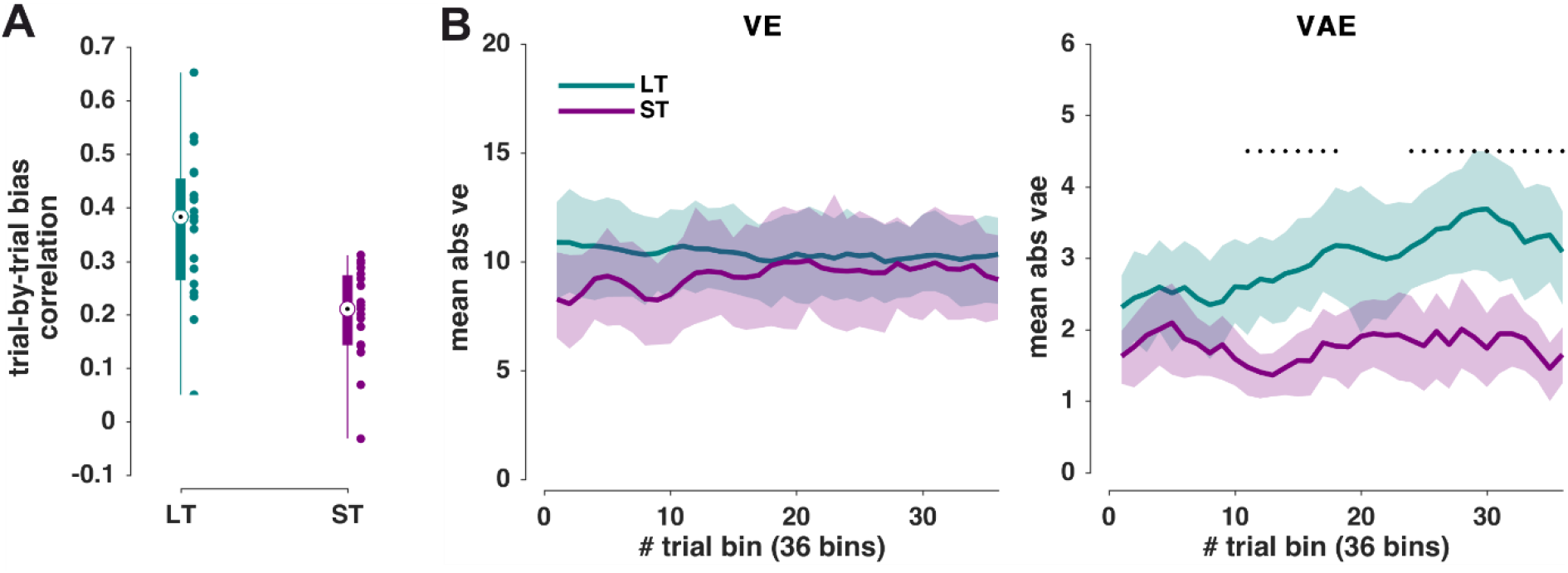
Trial-by-trial correlation of biases and temporal progression of biases. **(A)** Trial-by-trial correlation of *ve* and *vae* biases within participants. Boxplot and individual data (dots). The correlation was significant (p < 0.01, Spearman’s correlation) for 18 and 16 out of 19 participants for LT and ST. The median values for LT and ST were 0.382 and 0.211. Permutation test on the difference of medians confirmed that the correlation was stronger in the LT (p = 0.00016, permutation test based on 50000 random samples). **(B)** Temporal progression of biases. The LT data were averaged in increments of 5 trials resulting in 36 bins and the binned data were combined across blocks with leftward and rightward discrepancies. ST data were averaged in increments of 9 trials resulting in 36 bins. Shaded areas indicate 95% hybrid bootstrap confidence intervals. Black dots denote a significant difference between the LT and ST tested with a cluster-based permutation test (p < 0.01; See Methods for details). VE: ventriloquism effect. VAE: ventriloquism aftereffect.

**Figure S3.**
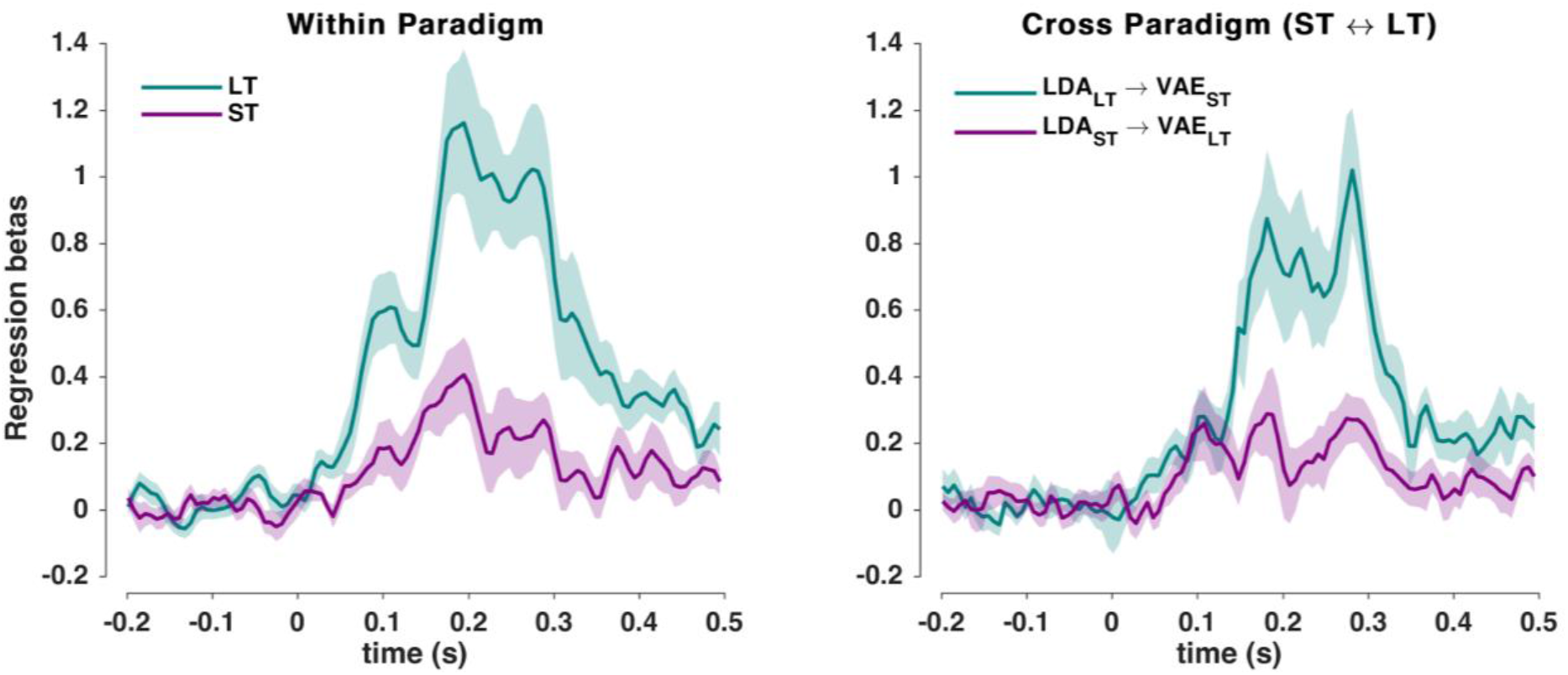
Time course of regression betas for the single-trial neuro-behavioral modeling of the aftereffect bias (c.f. Figure 2B,C). **(left)** Regression betas for the LDA_ΔVA from the within paradigm analyses. **(right)** Regression betas from the cross-paradigm analysis. Solid lines indicate the group-level mean, shaded areas are SEM across participants.

**Figure S4.**
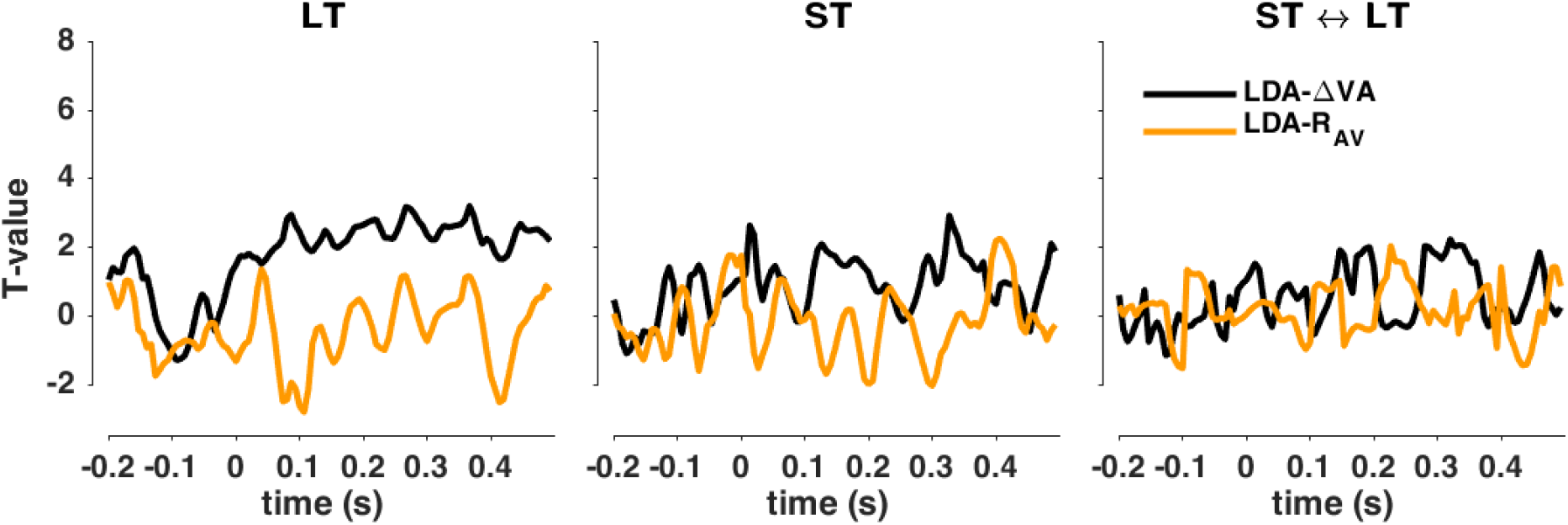
Control analysis using the EOG data to predict the ventriloquism aftereffect (analogous to Figure 2B,C). Classifying the direction of ΔVA from the EOG signals did not allow predicting the single trial bias in neither paradigm (left, center) nor across-paradigms (right). Graphs show group-level t-values. No significant clusters emerged at p<0.01.

